# Comparative analysis of single cell lung atlas of bat, cat, tiger and pangolin

**DOI:** 10.1101/2021.12.26.473325

**Authors:** Xiran Wang, Zhihua Ou, Peiwen Ding, Chengcheng Sun, Daxi Wang, Jiacheng Zhu, Wendi Wu, Yanan Wei, Xiangning Ding, Lihua Luo, Meiling Li, Wensheng Zhang, Xin Jin, Jian Sun, Huan Liu, Dongsheng Chen

## Abstract

Horseshoe bats (*Rhinolophus sinicus*) might help maintain coronaviruses severely affecting human health, such as SARS-CoV and SARS-CoV-2. It has long been suggested that bats may be more tolerant of viral infection than other mammals due to their unique immune system, but the exact mechanism remains to be fully explored. During the COVID-19 pandemic, multiple animal species were diseased by SARS-CoV-2 infection, especially in the respiratory system. Herein, single-cell transcriptomic data of the lungs of a horseshoe bat, a cat, a tiger, and a pangolin were generated. The receptor distribution of twenty-eight respiratory viruses belonging to fourteen viral families were characterized for the four species. Comparison on the immune-related transcripts further revealed limited cytokine activations in bats, which might explain the reason why bats experienced only mild diseases or even no symptoms upon virus infection. Our findings might increase our understanding of the immune background of horseshoe bats and their insensitivity to virus infections.

---

Bats are important reservoir hosts for a myriad of viruses including coronaviruses, rabies viruses, Hendra viruses, influenza viruses, etc. They were mostly asymptomatic or only developed mild diseases during viral infections by Ebola viruses, coronaviruses, henipaviruses, etc. The immune response induced by virus infection was shown to differ between human and bat cells^1,2^ and that bats may have their unique transcripts that are not present in other mammals^3^ Bats were found to have limited interferon activation due to mutation in the STING protein^4^ and have contracted type I IFNα locus but constitutive IFNα expression without viral stimulation^5^. The unique immune response pathways and antiviral gene expression profile of bats, may promote their tolerance to viral infections^6^. Bats are probably the initial host of SARS-CoV-2^7,8^, the etiological virus causing COVID-19. Although the direct progenitor of SARS-CoV-2 remains unknown, its closest relative (RaTG13) has been detected in a horseshoe bat (*Rhinolophus sinicus*), indicating horseshoe bats as its potential reservoir hosts. Moreover, horseshoe bats (genus *Rhinolophus*) were also found to harbor other groups of coronaviruses including the SARS-CoV emerged in China from 2002 to 2003^9–11^, indicating their critical role in the maintenance of human sensitive coronaviruses. Since the outbreak of COVID-19, multiple animal species have been infected and diseased by SARS-CoV-2, including pangolins, cats, tigers, etc.^12–16^. Herein, we conducted a comparative study using single cell transcriptomic data to elucidate the lung immune landscape of bat, cat, tiger and pangolin, which might help reveal the molecular basis for their differential immune behaviors upon infections by SARS-CoV-2.

Due to species-specific immune response upon viral infection, clinical symptoms in the lower respiratory differ among species. While bats, cats, tigers and pangolins were all permissive to SARS-CoV-2 infection, details of their biological background remain unknown. Herein, we collected lung tissues from healthy individuals of bats to generate single-nucleus libraries of lung cells, resulting in a total of 11838 pulmonary cells passing quality control (**Figures 1A and 1B**). Nine major cell types were identified in the lung atlas of bats, which included alveolar type 1 cells (AT1), alveolar type 2 cells (AT2), ciliated cells, secretory cells, endothelial cells, fibroblasts, T cells, B cells and macrophages, each demonstrating the specific expression of canonical cell type markers **(Figure 1C, Table S1**).

**Figure 1.**
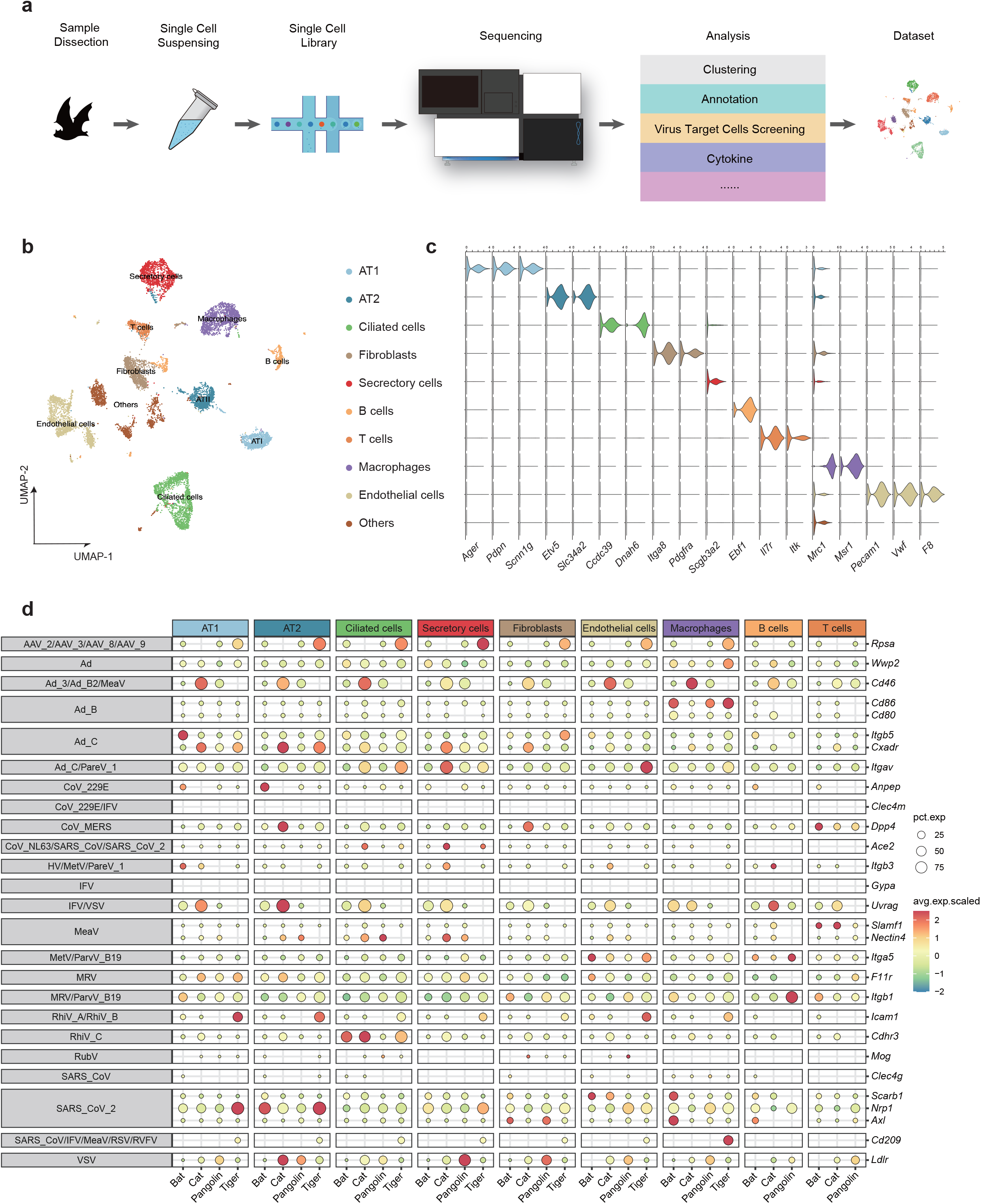
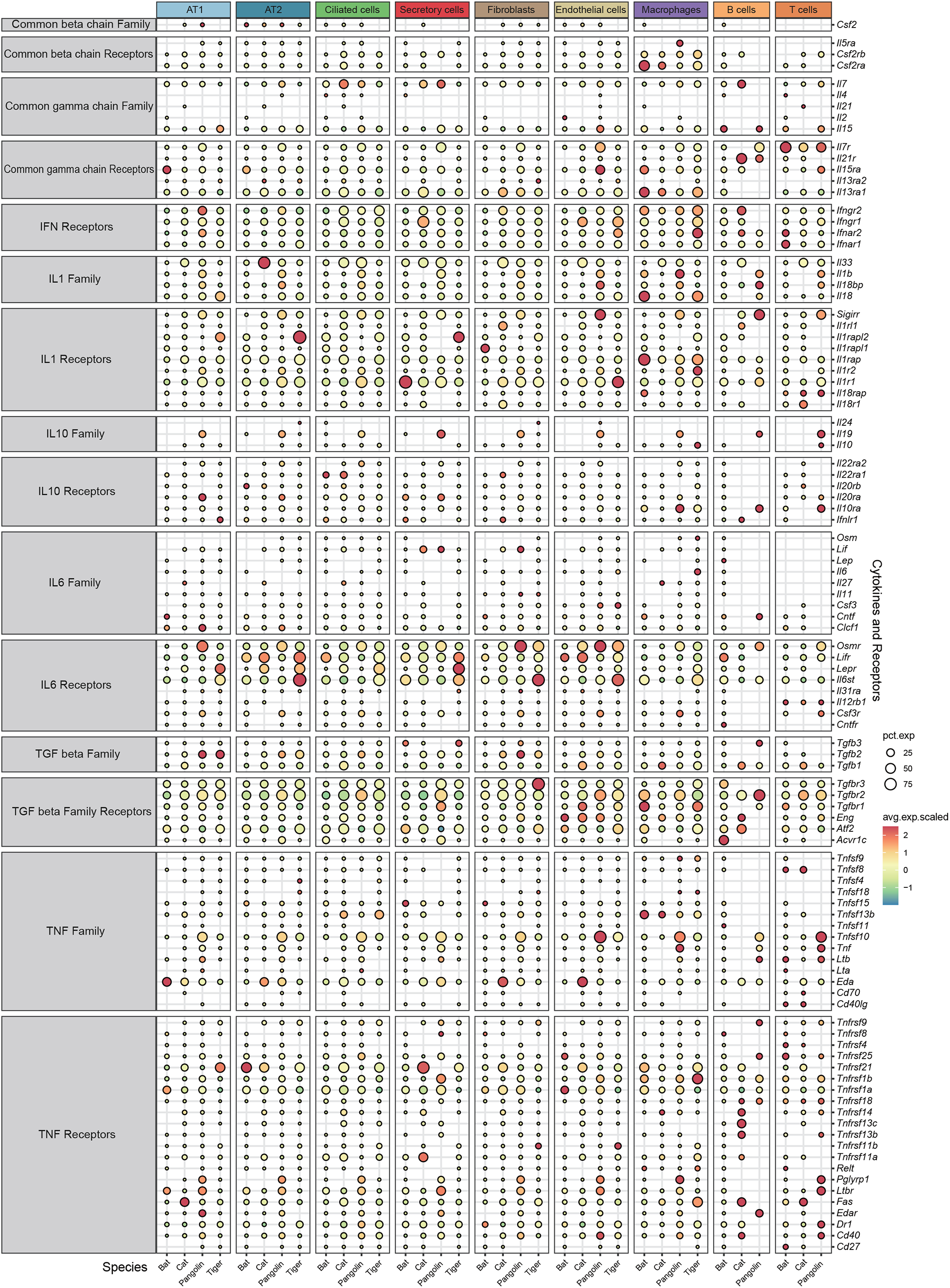
Comparative single cell lung atlas of the bat, cat, tiger and pangolin. **a**, Illustration of the overall project design. **b**, UMAP plot of bat lung. **c**. Violin plot showing the expression patterns of canonical cell type markers. **d**. Expression proportion and scaled expression value of virus receptors in distinct cell types of bat, cat, tiger and pangolin.

Because receptor binding is critical for viral entry into cells and that the distribution of receptors reveals the susceptibility of cells to viral infection, which may further stimulate the local immune response. Therefore, we determined the expression patterns of 29 genes encoding entry receptors of respiratory viruses in the lung cells of the four species. Compared with the other three species, *Itgb5* (a receptor of adenoviruses) and *Anpep* (a receptor of human coronavirus 229E) were enriched in bat AT1 and AT2, respectively. Another adenovirus receptor, *Cd86*, was enriched in the macrophages of bat, pangolin and tiger. The rhinovirus receptor, *Cdhr3*, was enriched in the ciliated cells of both bat and cat. Adeno-associated virus receptor *Rpsa* were significantly expressed in tiger pulmonary cells except unannotated T and B cells. *ACE2*, receptor for SARS-CoV and SARS-CoV-2, were relatively highly expressed in ciliated and secretory cells of cat and in the secretory cells of tiger. Only marginal expressions of *ACE2* were observed in bat ciliated cells. However, the receptor for SARS-CoV-2, *Scarb1*, displayed highly specific expression in bat endothelial cells and macrophages. Another two SARS-CoV-2 receptors, *Nrp1* and *Axl*, also showed significant cell type and species specificity. *Nrp1* was largely enriched in AT1/AT2 of tigers and AT2 of bats, whereas *Axl* was highly expressed in fibroblasts and macrophages of bat lung, and fibroblasts of pangolin lung (**Figure 1D**).

Cytokine storm, due to uncontrolled and excessive release of pro-inflammatory cytokines, is one of the main culprits contributing to severe lung pathogenesis caused by various virus infections^17^. As a natural reservoir for zoonotic viruses, bats display no significant symptoms after virus infection thanks to its unique immunity^18^. Here, we compared the expression profiles of a variety of pro-inflammatory cytokines (IL1, IL6, TNF, interferons) and anti-inflammation cytokines (IL10, TGF beta) among distinct pulmonary cell types of bat, cat, pangolin, and tiger. Regarding anti-inflammation cytokines and corresponding receptors, no significant differences were observed among the pulmonary cells of the four species (**Figure S1**). However, receptors for specific pro-inflammation cytokines, IL-6 (*Osmr, Lifr*) and interferons (IFN) (*Ifngr2, Ifnar1, Ifnar2*), were significantly lowly expressed in bat pulmonary cells (**Figure S1**). *Lif* was suggested to be a mediator of pro-inflammation in several inflammatory disorders^19^ and was lowly expressed in bat. Accumulating evidence has indicated that *OSM* mediates lung inflammations^20^ and that *Osmr* was abundantly expressed on mice pulmonary endothelial and fibroblast cells. Similarly, we observed enrichment of *Osmr* in these two cell types of cat and tiger, and wide expressions in all the nine pulmonary cell types of pangolin. Interestingly, expression of *Osmr* was almost lost in bat lung cells. Type 1 interferons (IFNs) play central roles in initiating lung inflammations^21^ and IFN-γ is a proinflammatory cytokine participated in inflammation and autoimmune disease^22^. Here, two receptors for type 1 interferon (*Ifnar1, Ifnar2*) and a receptor for IFN-γ (*Ifngr2*) were found to be depleted in bat cells (**Figure S1**). In summary, our results suggested that the transcriptions of some pro-inflammatory cytokine receptors were suppressed in bat lung cells, which could probably provide novel insights about the specific immune characteristics of bats.

Due to experimental limitations, we have only characterized the cytokine expressions of the lung cells of bats, cats, tiger and pangolins in healthy status, which may help unravel the baseline expression of immune factors. To fully understand immune responses stimulated by specific viral infections in bats, transcriptome data from appropriately controlled infection experiments are desired to better illustrate the differential pulmonary immune responses between these species. Moreover, because bats, tigers and pangolins were feral species, the animals sampled may not be strictly healthy as the pathogen-free laboratory animals. Therefore, our data might be influenced by individual bias to some extent.

In this study, we have generated the single-cell transcriptomes for the lungs of four non-model species associated with the cross-species transmissions of SARS-CoV-2. Besides, bats and animals such as cats and tigers are frequently involved in the cross-species transmissions of other viruses, such as various subtypes of influenza A viruses. Our transcriptome data revealed their cellular heterogeneity and thus laid the foundation for in-depth comparative study regarding the cellular and immune biology upon virus infections.

## MATERIAL AND METHODS

### Ethical statement

Sample collection and research were performed with the approval of Institutional Review Board on Ethics Committee of BGI (Approval letter reference number BGI-NO. BGI-IRB A20008). All procedures were conducted according to the guidelines of Institutional Review Board on Ethics Committee of BGI. The sampling procedures strictly followed the ‘Guidelines on Ethical Treatment of Experimental Animals’ established by the Ministry of Science and Technology, China.

### Sample collection and dissection

Bats used in this study were all male *Rhinolophus sinicus* (Chinese horseshoe bat) which were identified the species by the field experts, and obtained from Guangdong province, China, then dissected and stored in -80°C freezer immediately. After being isolated, the lung tissues of bats were rinsed by 1X PBS and stored in liquid nitrogen. Subsequently, we used mechanical extraction method from previous study^23^ to obtain the single nucleus and then stained using DAPI (4’,6-diamidino-2-phenylindole), calculated the density of the nucleus to ensure the quality of the single nucleus RNA sequencing library construction.

### Single cell RNA sequencing library preparation and sequencing

The isolated nuclei were separated from lung tissue of bats and then their reactions were performed according to the manufacturer’s protocol for the Chromium Single Cell 3’ GEM, Library & Gel Bead Kit v3.1. Library preparation was carried out following the guidelines provided by the manufacturer, total of 2 libraries were sequenced using a compatible Illumina NovaSeq 6000 platform.

### Processing of single-nucleus RNA-seq data

Raw sequencing data was aligned to ref genome sequence of *Rhinolophus sinicus* (GCF_001888835.1_ASM188883v1) and preprocessed by CellRanger 3.0.2 (10X Genomics). After obtaining the single cell gene expression matrices, we used Seurat 3.2.2^24^ to perform the downstream analysis. First, genes detected in less than three cells were discarded. Then, low-quality cells in which expressed gene numbers are less than 200 were filtered out. Moreover, cells with the percentage of mitochondrial genes more than 10% were removed. After quality control, “LogNormalize” function was used to normalize and “FindVariableGenes” was used to calculate the variance scores of each gene. Then, we applied “cca” to integrate the two libraries and remove the batch effect. Following, the integrated data was scaled and principal component analysis (PCA) was performed on the corrected data. Clusters were identified using “FindClusters” function and visualized by UMAP.

### Data collection

Other single cell RNA sequencing data (cat, tiger, pangolin) were obtained from public^1^.

### Annotation

The lung cell types of bats were annotated according to the expression of canonical markers. The lung cell types of cat, tiger and pangolin were acquired from Chen et al.

### DEGs identification and GO enrichment analysis

We applied “FindAllMarkers” function to identify the differentially expressed genes (DEGs). P value of the significance of DEGs were calculated by default Wilcox test and adjusted using Bonferroni methods. Genes of which adjusted p value is less than 0.05 and absolute value of log fold change is more than 0.25 were defined as DEGs and used for the following analysis. R package clusterProfiler^25^ was applied for GO term enrichment analysis.

### Cross-species data integration

To facilitate the comparation among four species, we first convert all genes of bats to the homogenous mouse genes using OrthoFinder^26^. Then we used Seurat^2^ to integrate the single cell datasets of bats, cat, tiger and pangolin.

### Cell type specific expression patterns of respiratory virus receptors and cytokine

Used the integrated datasets of four species, we calculated the average expression value and the percentage of expression for known respiratory virus receptors collected from previous study^27^ in various cell types. Besides, the expression patterns of cytokine genes in each cell type were also calculated using “DotPlot” function from Seurat and visualized by ggplot2^28^.

## ACKNOWLEDGMENTS

This work was supported by the Guangdong Major Project of Basic and Applied Basic Research (Grant 2020B0301030007), the Local Innovative and Research Teams Project of Guangdong Pearl River Talents Program (Grant 2019BT02N054), the Program for Changjiang Scholars and Innovative Research Team in University of Ministry of Education of China (Grant No. IRT_17R39), the Innovation Team Project of Guangdong University (Grant No. 2019KCXTD001)

## AUTHOR CONTRIBUTION

Conceptualization, D.C., J.S., H.L., J.Z., L.L. and X.D.; methodology, D.C., P.D., X.W., Z.O. and J.Z; software, X.W. and P.D.; validation, X.W. P.D. and Z.O.; formal analysis, X.W. and P.D.; investigation, C.S., X.W., P.D., W.W. and Y.W.; resources, M.L. D.C. and X.J.; data curation, X.W. and P.D.; writing - original draft preparation, X.W., Z.O. and P.D.; writing - review and editing, X.W., Z.O., P.D., J.S., W.Z., and D.C.; visualization, P.D. and X.W.; supervision, J.S., H.L. and D.C.; project administration, D.W., J.S., H.L. and D.C.; funding acquisition, J.S. D.C. and H.L.. All authors have read and agreed to the published version of the manuscript.

## DATA AVAILABILITY STATEMENT

The raw data supporting the findings of this study will be made available upon request. The data that support the findings of this study have been deposited into CNGB Sequence Archive (CNSA)^29^ of China National GeneBank DataBase (CNGBdb)^30^ with accession number CNP0002166.

## SUPPLEMENTARY FIGURES AND TABLES

**Figure S1: Dot plot showing the expression percentage and scaled expression value of cytokine and their receptors in distinct cell types of bat, cat, tiger and pangolin**.

**Table S1: Differentially expressed genes (DEGs) of bat lung**.

